# Rare schizophrenia risk variants are enriched in genes shared with neurodevelopmental disorders

**DOI:** 10.1101/069344

**Authors:** Tarjinder Singh, James T. R. Walters, Mandy Johnstone, David Curtis, Jaana Suvisaari, Minna Torniainen, Elliott Rees, Conrad Iyegbe, Douglas Blackwood, Andrew M. McIntosh, Georg Kirov, Daniel Geschwind, Robin M. Murray, Marta Di Forti, Elvira Bramon, INTERVAL Study, UK10K Consortium, Aarno Palotie, Michael C. O’Donovan, Michael J. Owen, Jeffrey C. Barrett

## Abstract

By meta-analyzing rare coding variants in whole-exome sequences of 4,264 schizophrenia cases and 9,343 controls, *de novo* mutations in 1,077 trios, and array-based copy number variant calls from 6,882 cases and 11,255 controls, we show that individuals with schizophrenia carry a significant burden of rare damaging variants in a subset of 3,230 “highly constrained” genes previously identified as having near-complete depletion of protein truncating variants. Furthermore, rare variant enrichment analyses demonstrate that this burden is concentrated in known autism spectrum disorder risk genes, genes diagnostic of severe developmental disorders, and the autism-implicated sets of promoter targets of *CHD8*, and mRNA targets of *FMRP*. We further show that schizophrenia patients with intellectual disability have a greater enrichment of rare damaging variants in highly constrained genes and developmental disorder genes, but that a weaker but significant enrichment exists throughout the larger schizophrenia population. Combined, our results demonstrate that schizophrenia risk loci of large effect across a range of variant types implicate a common set of genes shared with broader neurodevelopmental disorders, suggesting a path forward in identifying additional risk genes in psychiatric disorders and further supporting a neurodevelopmental etiology to the pathogenesis of schizophrenia.

## Introduction

Schizophrenia is a common and debilitating psychiatric illness characterized by positive symptoms (hallucinations, delusions, disorganized speech and behaviour) and negative symptoms (social withdrawal and diminished emotional expression) that result in social and occupational dysfunction^1,2^. Operational diagnostic criteria for the disorder as described in the DSM-V require the presence of at least two of the core symptoms over a period of six months with at least one month of active symptoms^3^. It is widely recognized that current categorical psychiatric classifications have a number of shortcomings, in particular that they overlook the increasing evidence for etiological and mechanistic overlap between psychiatric disorders^4^.

It seems likely that a diverse range of pathophysiological processes contribute to the clinical features of schizophrenia^5^. Indeed, previous studies have suggested a number of hypotheses about schizophrenia pathogenesis, including abnormal pre-synaptic dopaminergic activity^6^, postsynaptic mechanisms involved in synaptic plasticity^7^, dysregulation of synaptic pruning^8^, and disruption to early brain development^9,10^. This postulated pathophysiological complexity is underpinned by the varied nature of genetic contributions to risk of schizophrenia. Genome-wide association studies have identified associations to over 100 independent loci defined by common (minor allele frequency [MAF] > 2%) single nucleotide variants (SNVs) ^11^, and a recent analysis determined that more than 71% of all one-megabase regions in the genome contain at least one common risk allele^12^. The modest effects of these variants (median odds ratio [OR] = 1.08) combine to produce a polygenic contribution that explains only a small fraction 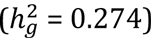 of the overall liability^12^. In addition, a number of rare variants have been identified that have very large effects on risk. These are best exemplified by eleven large, rare recurrent copy number variants (CNVs) but evidence from whole-exome sequencing studies implies that many other rare coding SNVs and *de novo* mutations also confer substantial risk^13-16^. There is growing evidence that some of the same genes and pathways are affected by both common and rare variants^7,17^. Pathway analyses of common variants and hypothesis-driven gene set analyses of rare variants have begun to enumerate some of these specific biological processes, including histone methylation, transmission at glutamatergic synapses, calcium channel signaling, synaptic plasticity, and translational regulation by the fragile × mental retardation protein (*FMRP*)^11,18,19^.

In addition to exploring the biological mechanisms underlying schizophrenia, genetic analyses can also be used to understand its relationship to other neuropsychiatric and neurodevelopmental disorders. For example, schizophrenia, bipolar disorder, and autism (ASD) show substantial sharing of common risk variants^20,21^. Sequencing studies of neurodevelopmental disorders suggest that this sharing of genetic risk may extend to rare variants of large effect. In the largest sequencing study of ASD to date, 20 of the 46 genes and all six CNVs implicated (false discovery rate [FDR] < 5%) had been previously described as dominant causes of developmental disorders^22^. Furthermore, *de novo* loss-of-function (LoF) mutations identified in ASD probands in these genes were disproportionately found in individuals with cognitive impairment^22,23^. The evidence from rare variants for a broader shared genetic etiology between schizophrenia and developmental disorders is more mixed, with a nominal overlap of *de novo* mutations^13^, but no evidence in a whole-exome sequencing study of 2,536 cases and 2,543 controls^24^. Intriguingly, the 11 rare schizophrenia CNVs described above also increase risk for intellectual disability and other congenital defects^16,25^, and more recently, a meta-analysis of whole-exome sequence data showed that LoF variants in *SETD1A* confer substantial risk for both schizophrenia and developmental disorders^17^. Emerging results from these individual risk loci showing pleiotropic effects offers the possibility that a larger number of developmental disorder genes could additionally confer substantial risk for schizophrenia. Here, we present a detailed analysis of the largest accumulation of schizophrenia rare variant data assembled to date, in order to better understand which genes are implicated by these variants, and how they relate to neurodevelopment more generally.

## Results

### Study design

To maximize our power to detect signals of enrichment of damaging variants in schizophrenia cases in groups of genes, we performed a meta analysis of three different types of rare coding variant studies. We first incorporated high-quality SNV calls from whole-exome sequences of 4,264 schizophrenia cases and 9,343 matched controls, then aggregated *de novo* mutations identified in 1,077 schizophrenia parent-proband trios (Figure 1), and finally added CNV calls from genotyping array data of 6,882 cases and 11,255 controls. The ascertainment of these samples, data production, and quality control were described previously^17,26^. All *de novo* mutations included in our analysis had been validated through Sanger sequencing, and stringent quality control steps were performed on the case-control data to ensure that sample ancestry and batch were closely matched between cases and controls (Online Methods).

**Figure 1:**
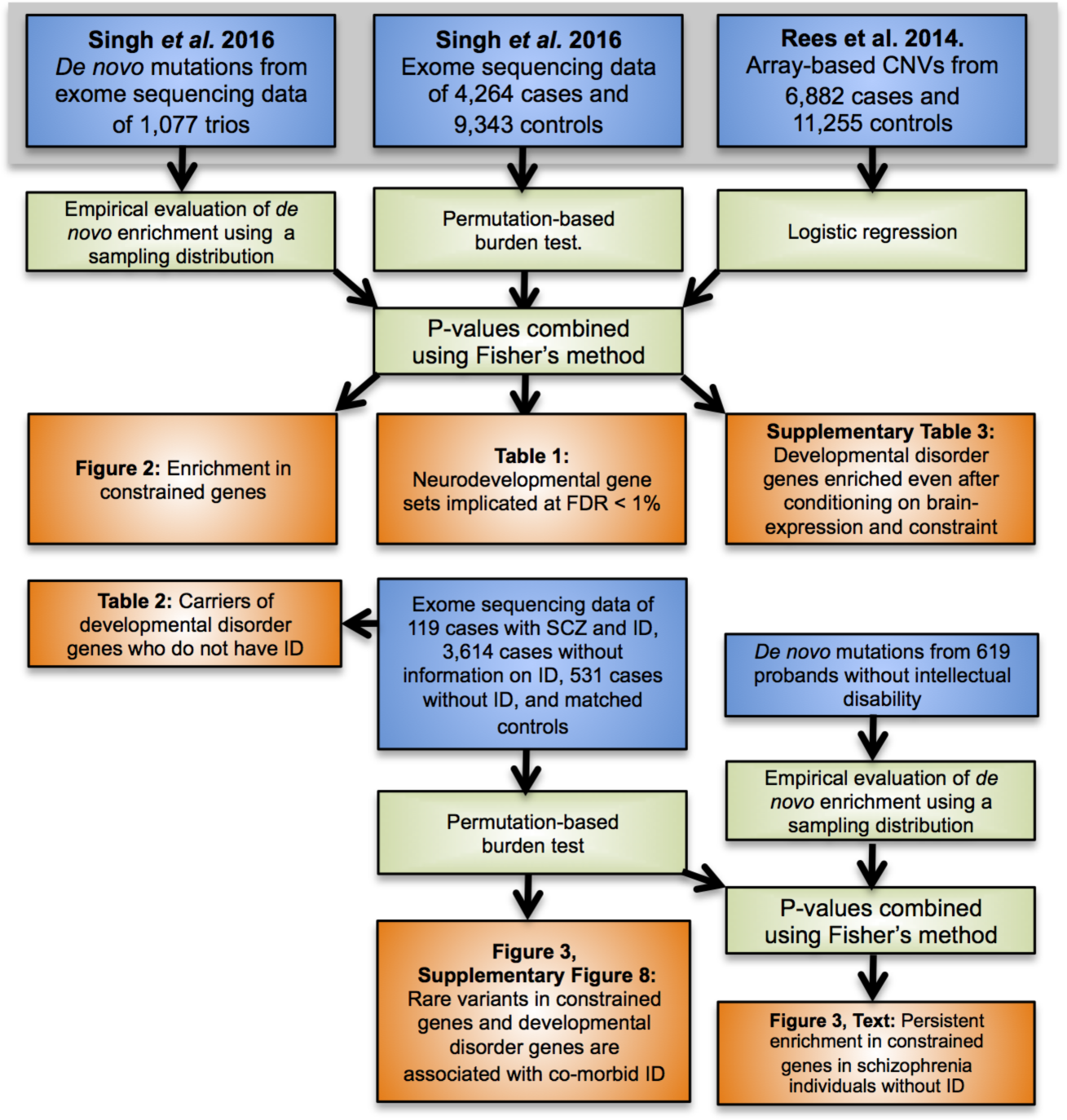
Analysis workflow. Data sets are shown in blue, statistical methods and analysis steps are shown in green, and results (figures and tables) from the analysis are shown in orange.

For each data type, we used previously described methods appropriate to those data to test for an excess of rare variants (Figure 1, Online Methods). For case-control SNV data, we performed permutation-based gene set burden tests corrected for exome-wide differences between cases and controls^17,24^. For case-control CNV data, we used a logistic regression framework that compares the rate of CNVs overlapping a specific gene set while correcting for differences in CNV size and number of genes disrupted^7,18,27^. Lastly, we compared a sampling distribution of *de novo* mutations based on a gene-specific mutation rate model to the observed number of mutations in each gene set to calculate an empirical *P*-value. In all three analyses, we chose thresholds for consequence severity and minor allele frequency that showed the strongest enrichment in previously implicated schizophrenia gene sets (Supplementary Figure 1, Online Methods). We restricted our analysis to case-control LoF variants observed only once in our data set and absent in the ExAC database^28^, small deletions and duplications overlapping fewer than seven genes with MAF < 0.1% (Supplementary Figure 2), and *de novo* mutations annotated as LoF or damaging missense (CADD phred score ≥ 15)^29^.

We tested for an excess of rare damaging variants in schizophrenia patients in 1,776 gene sets (Online Methods, Supplementary Table 1, and detailed results below). Gene set *P*-values were computed using the three methods and variant definitions described above, and then meta-analyzed using Fisher’s Method to provide a single *P*-value for each gene set. Because we gave each data type equal weights, gene sets achieving significance typically show at least some signal in all three types of data. We observed a marked inflation in the quantile-quantile (Q-Q) plot of gene set *P*-values (Supplementary Figure 3), so we conducted two analyses to ensure our results were robust and not biased due to methodological or technical artifacts in our data. First, we observed no inflation of *P*-values when testing for enrichment of synonymous variants in our case-control and *de novo* analyses (Supplementary Figure 3). Second, we created random gene sets by sampling uniformly across the genome, and observed null distributions in Q-Q plots regardless of variant class and analytical method (Supplementary Figure 4). These findings suggested that our methods sufficiently corrected for known genome-wide differences in LoF and CNV burden between cases and controls, and other technical confounders like batch and ancestry.

### Rare, damaging schizophrenia variants are concentrated in constrained genes

A recent study calculated gene-level selective constraint for every gene in the genome by comparing the observed number of rare protein-truncating SNVs in exomes from 60,706 individuals without severe, early-onset disorders to the number predicted from a gene-specific mutation rate model^30^. They identified 3,230 genes with near-complete depletion of such truncating variants^28^, which we refer to as the “highly constrained” gene set. Notably, recurrent *de novo* LoF mutations identified in individuals with ASD or developmental disorders are overwhelmingly concentrated in this set of genes^22,28,30^, suggesting that at least some of the constraint is driven by severe neurodevelopmental consequences of having only one functioning copy of these genes.

We found that rare damaging variants in schizophrenia cases were also enriched in the highly constrained gene set (P < 1×10^−14^, Table 1, Figure 2), with support in case-control SNVs (P < 5×10^−7^; OR 1.21, 1.12-1.30, 95% CI), case-control CNVs (P = 2.6×10^−4^; OR 1.22, 1.15 - 1.30, 95% CI), and *de novo* mutations (P = 1×10^−6^; OR 1.34, 1.2 - 1.48, 95% CI). While this result was consistent with observations in intellectual disability and ASD^30,31^ the absolute effect size is much smaller (e.g. *de novos*, Supplementary Figure 5 and 6). We observed no excess burden of rare damaging variants in the remaining 14,996 genes (Figure 2, Supplementary Figure 6 and 7). Furthermore, this signal was spread among many different constrained genes: if we rank genes by decreasing significance, the enrichment disappears in the case-control SNV analysis (P > 0.05) only after the exclusion of the top 50 genes. This suggests that the contribution of damaging rare variants in schizophrenia is not concentrated in just a handful of genes, but instead spread across many genes.

**Table 1:** Gene sets enriched for rare coding variants conferring risk for schizophrenia at FDR < 1%. The effect sizes and corresponding *P*-values from enrichment tests of each variant type (case-control SNVs, DNM, and case-control CNVs) are shown for each gene set, along with the Fisher’s combined P-value (P_meta_) and the FDR-corrected Q-value (Q_meta_). We only show the most significant gene set if there are multiple ones from the same data set or biological process (see Supplementary Table 1 for all 1,776 gene sets). All gene sets displayed had been previously implicated in ASD and ID. N_genes_: number of genes in the gene set; Est: effect size estimate and its lower and upper bound assuming a 95% CI; DNM: *de novo* mutations.

| Gene set | N<sub>genes</sub> | Est<sub>SNV</sub> | P<sub>SNV</sub> | Est<sub>DNM</sub> | P<sub>DNM</sub> | Est<sub>CNV</sub> | P<sub>CNV</sub> | P<sub>meta</sub> | Q<sub>meta</sub> |
| --- | --- | --- | --- | --- | --- | --- | --- | --- | --- |
| ExAC constrained genes (pLI > 0.9) | 3230 | 1.21 (1.12-1.3) | < 5.00 x 10<sup>-7</sup> | 1.34 (1.2-1.48) | 1.00 x 10<sup>-6</sup> | 1.22 (1.15-1.3) | 0.00026 | <9.00 x 10<sup>-14</sup> | <1.60 x 10<sup>-10</sup> |
| Sanders et al. autism risk genes (FDR < 10%) | 66 | 1.5 (1.06-2.11) | 0.012 | 2.16 (1.17-4.06) | 0.026 | 2.21 (1.75-2.79) | 0.00033 | 1.60 x 10<sup>-5</sup> | 0.0046 |
| Dominant, diagnostic DDG2P genes, in which LoF variants result in developmental disorders with brain abnormalities | 125 | 1.82 (1.22-2.69) | 0.0046 | 2.17 (1.32-3.51) | 0.0047 | 1.67 (1.33-2.11) | 0.012 | 3.50 x 10<sup>-5</sup> | 0.0061 |
| Darnell et al. targets of FMRP | 790 | 1.29 (1.15-1.44) | 1.20 x 10<sup>-5</sup> | 1.57 (1.29-1.89) | 2.30 x 10<sup>-5</sup> | 1.32 (1.2-1.47) | 0.0032 | 3.70 x 10<sup>-10</sup> | 2.20 x 10<sup>-7</sup> |
| Cotney et al. CHD8-targeted promoters (hNSC and human brain tissue) | 2920 | 1.11 (1.04-1.19) | 0.00094 | 1.38 (1.18-1.6) | 0.00011 | 1.11 (1.05-1.18) | 0.027 | 5.70 x 10<sup>-7</sup> | 0.0002 |
| Cerebellum-expressed genes (Brainspan) | 15976 | 1.03 (1-1.06) | 0.012 | 1.05 (1.01-1.08) | 0.015 | 1.11 (1.07-1.14) | 0.0011 | 2.50 x 10<sup>-5</sup> | 0.0055 |
| GOCC: chromatin (GO:0000785) | 429 | 1.32 (1.08-1.6) | 0.0007 | 2.01 (1.39-2.89) | 0.00076 | 0.76 (0.67-0.88) | 0.98 | 6.30 x 10<sup>-5</sup> | 0.0085 |

**Figure 2:**
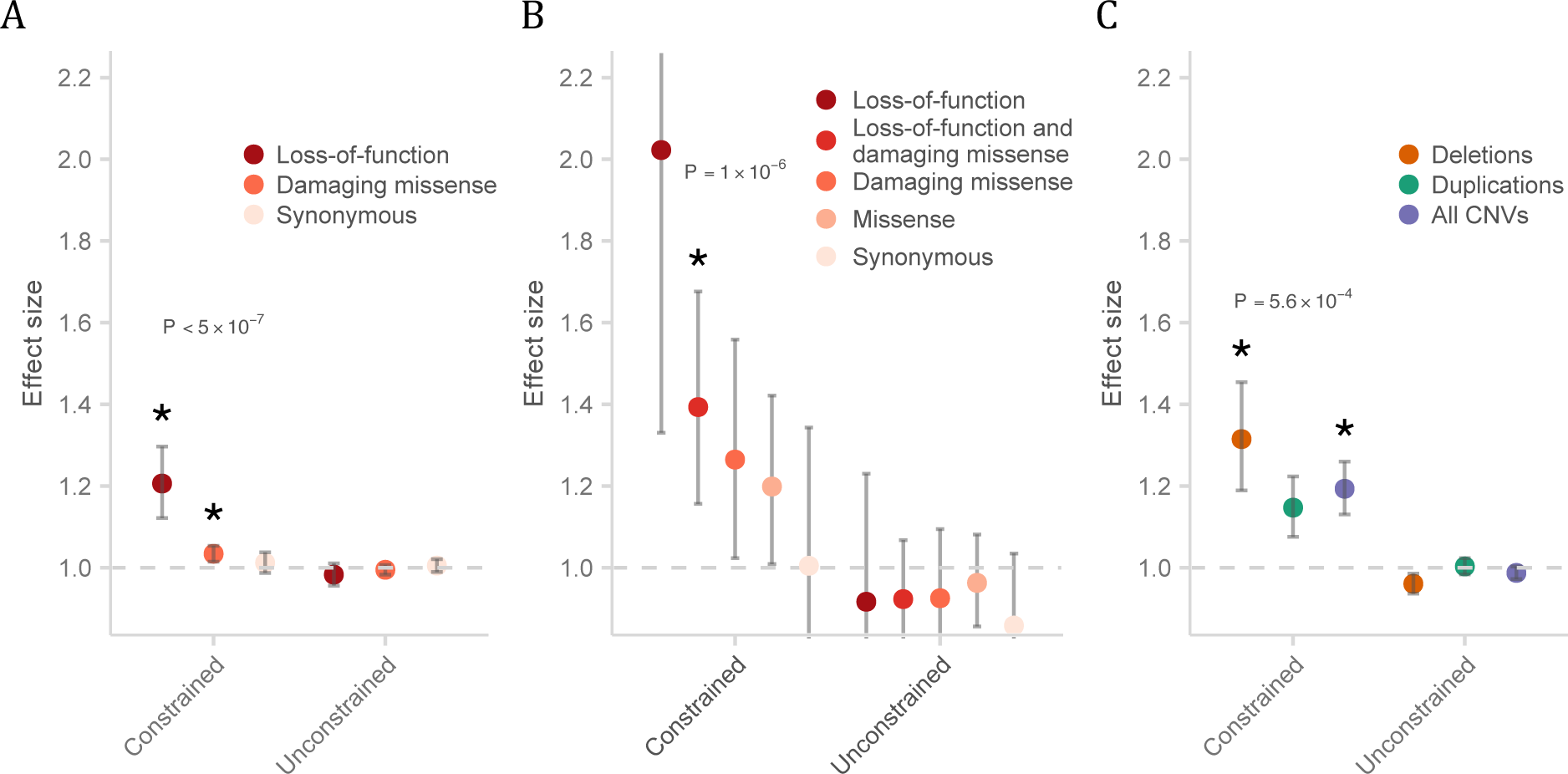
Enrichment of schizophrenia rare variants in constrained genes. **A:** Schizophrenia cases compared to controls for rare SNVs and indels; **B:** Rates of *de novo* mutations in schizophrenia probands compared to control probands; **C:** Case-control CNVs. *P*-values shown were from the test of LoF enrichment in **A**, LoF and damaging missense enrichment in **B**, and all CNVs enrichment in **C**. Error bars represent the 95% CI of the point estimate. Constrained: 3,230 genes with near-complete depletion of truncating variants in the ExAC database; Unconstrained: genes not under genic constraint; Damaging missense: missense variants with CADD phred > 15. Asterisk: P < 1 × 10^−3^.

### Schizophrenia risk genes are shared with other neurodevelopmental disorders

Given the significant enrichment of rare damaging variants in constrained genes in DD, ASD and schizophrenia, we next asked whether these variants affected the same genes. We found that both ASD risk genes identified from exome sequencing meta-analyses^22^ and genes in which LoF variants are known causes of severe developmental disorders as defined by the DDD study^32^ were significantly enriched for rare variants in individuals with schizophrenia (*P*_ASD_ = 1.6×10^−5^; *P*_DD_ = 3.5×10^−5^; Table 1, Online Methods). Previous analyses have shown an enrichment of rare damaging variants in mRNA targets of *FMRP* in both schizophrenia and autism^13,24,31^, so we sought to identify further shared biology by testing targets of neural regulatory genes previously implicated in autism^31,33^. We observed enrichment of both such sets: promoter targets of *CHD8* (*P* = 5.7×10^−7^) and splice targets of *RBFOX* (*P* = 3.3×10^−4^) (Table 1).

We also tested an additional 1,769 gene sets from databases of biological pathways with at least 100 genes, as we lacked power to detect weak enrichments in smaller sets (Online Methods). We observed enrichment of damaging rare variants in schizophrenia patients at FDR *q* < 0.05 in 62 of these sets (Supplementary Table 1, 2). These included both previously implicated gene sets, like calcium signaling genes and glutamatergic synaptic density proteins comprising the NMDAR and ARC complexes^13,14,24,34^, as well as novel gene sets, such as regulation of ion transmembrane transport (GO:0034765) and neuron projection morphogenesis (GO:0048812). Notably, the gene sets most significantly enriched (FDR *q* < 0.01) for schizophrenia rare variants (Table 1) were all neurodevelopmental gene sets previously implicated in autism and intellectual disability (mRNA targets of *FMRP*, chromatin-associated genes [GO], promoter targets of *CHD8*^31-34^) as well as a large and generic set of cerebellum expressed genes. These FDR < 1% neurodevelopmental gene sets were significant even after conditioning on brain expression (Supplementary Tables 3, Online Methods), suggesting they represent more specific biological processes involved in schizophrenia. Notably, only known ASD risk genes (*P* = 1.6×10^−4^) and diagnostic DD genes (*P* = 1.2 x10^−3^) had an excess of rare coding variants above the enrichment already observed in constrained genes (Supplementary Table 3). Thus, in addition to biological pathways implicated specifically in schizophrenia, at least a portion of the schizophrenia risk conferred by rare variants of large effect is shared with childhood onset disorders of neurodevelopment.

### Schizophrenia rare variants are associated with intellectual disability

In the autism spectrum disorders, the observed excess of rare damaging variants has been shown to be much greater in individuals with intellectual disability than those with normal levels of cognitive function^23^. We observed a similar reduction in cognitive function in schizophrenia patients carrying *SETD1A* LoF variants^17^, so next sought to explore whether this pattern is consistent in schizophrenia in a wider set of genes. Our exome sequenced cases included 119 individuals specifically recruited because they had pre-morbid intellectual disability (IQ between 50 and 75) in addition to fulfilling the full diagnostic criteria for schizophrenia (Online Methods. We also aggregated relevant cognitive phenotype data for 697 additional cases, and identified a set of 531 whom we could confirm do not have intellectual disability (after excluding pre-morbid IQ < 85, fewer than 14 years of schooling or lowest decile of composite cognitive measures, depending on available data, Online Methods. Finally, we were left with 3,614 cases without any data on intellectual disability. When stratifying into these three groups (intellectual disability, unknown, no intellectual disability), we observed that the burden of damaging rare variants in constrained genes is significantly greater in the small set of cases with confirmed intellectual disability than in the remaining schizophrenia cases and controls (Figure 3). We also discovered that these individuals had a significantly elevated number of variants in diagnostic developmental disorder genes compared to the remaining cases and controls (Supplementary Figure 8), and identified carriers of LoF variants in *KMT2A* and *KMT2D* (the same family of lysine methyltransferases as *SETD1A*, also known as *KMT2F*).

**Figure 3:**
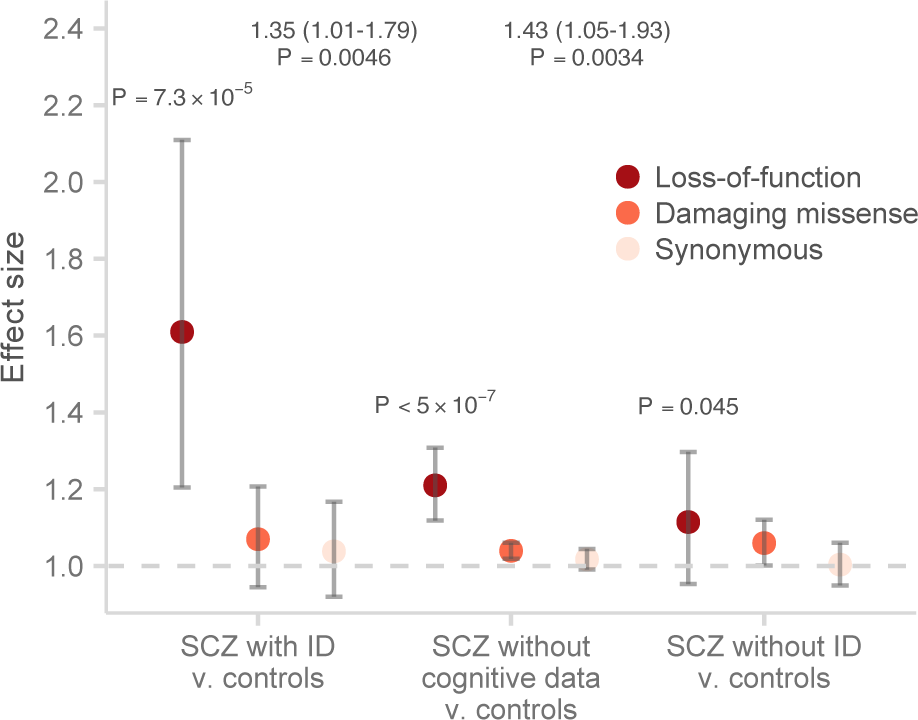
Enrichment of rare variants in constrained genes between schizophrenia (SCZ) individuals with ID, schizophrenia individuals without cognitive data, schizophrenia individuals without ID, and matched controls. The *P*-values shown were calculated from the burden test of LoF variants between the corresponding cases and matched controls. The enrichment of LoF variants in constrained genes between SCZ individuals with ID and other two SCZ groups (without cognitive data and without ID) were displayed as effect sizes and *P*-values above the case-control comparisons. Error bars represent the 95% CI of the point estimate. Damaging missense: missense variants with CADD phred > 15.

While the damaging rare variants in constrained genes were most highly enriched in the subset of schizophrenia patients with intellectual disability, we still observed a significant enrichment in the individuals whose intellectual disability status is unknown (*P* < 5×10^−7^, Figure 3), most of whom we expect to not have intellectual disability (Online Methods). Rare variants in constrained genes were enriched even in the set of individuals for whom intellectual disability can be excluded, and this result substantially increases in significance if we meta-analyze it with LoF and damaging missense de novo mutations from individuals without intellectual disability (*P*_case-control_ = 0.045, *P*_de novo_ = 4× 10^−4^, *P*_meta_ = 3.2×10^−4^, Online Methods). We additionally found twelve schizophrenia cases without ID carrying LoF variants in genes in which these variants are known causes of severe developmental disorders, supporting a nominal signal in this gene set (P_case-control_ = 0.069, *P*_de novo_ = 0.025, *P*_meta_ = 0.015). These individuals had symptoms satisfying the full diagnostic criteria for schizophrenia without signs of pre-morbid cognitive impairment (Table 2). Thus, rare damaging variants in constrained genes in individuals with schizophrenia follow the pattern previously described in autism: concentrated in individuals with intellectual disability, but are not exclusive to that group.

**Table 2:** Phenotypes of schizophrenia individuals with cognitive information carrying LoF variants in developmental disorder genes. Of the 531 UK10K schizophrenia individuals without intellectual disability, we acquired detailed clinical information for four out of the eight carriers of LoF variants in severe developmental disorders genes. These variants were observed only once in our data set and absent in the ExAC database. For each LoF variant, we provide its genomic coordinates (hg19) and the gene disrupted, the number of high-quality LoF variants within this gene identified in 60,706 ExAC individuals and the corresponding pLI score, and the expected developmental disorder syndrome according to DECIPHER. For each carrier, we describe notable neuropsychiatric symptoms (Clinical features), the level of education achieved (Education attainment), and the predicted pre-morbid IQ as extrapolated from National Adult Reading Test (NART). These four carriers satisfy the full diagnostic criteria for schizophrenia, and do not appear to be outliers in the expected cognitive range of schizophrenia patients. To identify high-quality ExAC LoF variants, we retained only variants in the canonical transcript and were called as homozygote (and not missing) in at least 85% of the ExAC data set (accessed on July 4^th^, 2016).

| Variant | Gene | nLoF<br>in<br>ExAC | pLI | Expected syndrome<br>based on DDG2P | Clinical features | Educational<br>attainment | Predicted<br>pre-<br>morbidity IQ |
| --- | --- | --- | --- | --- | --- | --- | --- |
| 1:151377686_GA/G<br>frameshift variant | POGZ | 2 | 1.000 | Intellectual Disability | Acute onset at age 20, treatment-resistant schizophrenia with severe depressive and negative symptoms. | Attended mainstream school, and achieved A level exams. | 105 |
| 1:151400550_C/T<br>stop gained | POGZ | 2 | 1.000 | Intellectual Disability | Paranoid schizophrenia, moderate negative symptoms, alcohol dependence. | Attended mainstream school, and achieved A level exams. | 108 |
| 11:102076807_T/C<br>splice donor variant | YAP1 | 0 | 0.999 | Coloboma, ocular,<br>with or without<br>hearing impairment,<br>cleft lip/palate,<br>and/or mental<br>retardation | Schizoaffective depression diagnosis,<br>prominent negative symptoms. | Attended mainstream school, but left with no qualification. | 86 |
| 16:89367335_C/A<br>stop gained | ANKRD11 | 2 | 1.000 | KBG syndrome | Late age of onset at age > 35, severe psychosis with first rank symptoms, multiple admissions, marked negative symptoms, and depression. | Attended mainstream school, but left with no qualification. | 102 |

## Discussion

Our integrated analysis of rare variants from thousands of whole-exome sequences provides evidence for a shared genetic etiology between schizophrenia and other neurodevelopmental disorders. While the identification of individual genes remains difficult at current samples sizes, we show that the burden of damaging *de novo* mutations, rare SNVs and CNVs in schizophrenia is primarily concentrated in the highly constrained set of 3,230 genes, an observation shared with autism and intellectual disability. Enrichment analyses in 1,776 gene sets further demonstrate that the most robust burden of rare variants in schizophrenia resides in genes in which LoF variants are diagnostic for severe developmental disorders and in known autism risk genes. In so far as the genes responsible for intellectual disability necessarily have effects during central nervous system development, and those that influence ASD must exert their effects in infancy at the very latest, the findings demonstrate that genetic perturbations adversely affecting nervous system development also increase risk for schizophrenia. Our findings therefore suggest that severe, psychiatric illnesses manifesting in adulthood can have origins early in development.

We additionally show that some of these perturbations have clear manifestations in childhood, and that risk variants of large effect in schizophrenia are particularly associated with pre-morbid intellectual disability. Our observations are consistent with results in autism in which recurrent *de novo* events are associated with cognitive impairment^21-23^. A weaker but still significant rare variant burden was observed in schizophrenia patients without intellectual disability, showing that variants of large effect do not simply confer risk for a small subset of schizophrenia patients but are relevant to disease pathogenesis more broadly.

Our data support the general observation that genetic risk factors for psychiatric and neurodevelopmental disorders do not follow clear diagnostic boundaries, and that the variants disrupting the same genes, and quite possibly, the same biological processes, result in a large range of phenotypic manifestation. For instance, a number of schizophrenia patients without intellectual disability carry LoF variants in developmental disorder genes. This clinically variable presentation is reminiscent of LoF variants in *SETD1A* and 11 large copy number variant syndromes, previously shown to confer risk for schizophrenia in addition to other prominent developmental defects^16,17^. We do not preclude the possibility that allelic series of LoF variants exist in these genes; however, the most common deletion in the 22q11.2 locus and a recurrent two base deletion in *SETD1A* are associated with both schizophrenia and more severe neurodevelopmental disorders, suggesting the same variants confer risk for a range of clinical features^17,35,36^. Ultimately, it may prove difficult to clearly partition patients genetically into subtypes with similar clinical features, especially if genes and variants previously thought to cause well-characterized Mendelian disorders can have such varied outcomes. This pattern is consistent with the hypothesis that LoF variants in constrained genes result in a spectrum of neurodevelopmental outcomes with the burden of mutations highest in intellectual disability and least in schizophrenia, corresponding to a gradient of neurodevelopmental pathology indexed by cognitive impairment^4^.

Despite the complex nature of genetic contributions to risk of schizophrenia, it is notable that across study design (trio or case-control) and variant class (SNVs or CNVs), risk loci of large effect are concentrated in a small subset of genes. Previous rare variant analyses in other neurodevelopmental disorders, such as autism, have successfully integrated information across *de novo* SNVs and CNVs to identify novel risk loci^22^. As sample sizes increase, meta-analyses leveraging the shared genetic risk across study designs and variant types will be similarly well powered to identify additional risk genes in schizophrenia.

## Author contributions

T.S., A.P., M.C.O., M.J.O., J.C.B conceived and designed the experiments.

T.S performed the statistical analysis.

T.S., J.T.R.W., M.J., D.C., J.S., M.T., E.R analysed the data.

T.S., J.T.R.W., M.J., J.S., M.T., E.R., C.I., D.B., A.M.M., G.K., D.G., R.M.M., M.D.F., A.P.,

M.C.O., M.J.O., J.C.B contributed reagents/materials/analysis tools.

T.S., D.C., M.C.O., M.J.O., J.C.B wrote the paper.

## Acknowledgements

We thank the thousands of patients who participated in these studies. We thank Timi Touloupoulou, Marco Picchioni, Chiara Nosarti, Fiona Gaughran, and Oliver Howes for contributing clinical data used in this study. The UK10K project was funded by Wellcome Trust grant WT091310. The INTERVAL sequencing studies are funded by Wellcome Trust grant WT098051. T.S. is supported by the Williams College Dr. Herchel Smith Fellowship. A.P. is supported by Academy of Finland grants 251704 and 286500, NIMH U01MH105666 and the Sigrid Juselius Foundation. The work at Cardiff University was funded by Medical Research Council (MRC Centre (G0801418 and Program Grants (G0800509. Participants in INTERVAL were recruited with the active collaboration of NHS Blood and Transplant England, which has supported fieldwork and other elements of the trial. DNA extraction and genotyping was funded by the National Institute of Health Research (NIHR), the NIHR BioResource and the NIHR Cambridge Biomedical Research Centre. The academic coordinating centre for INTERVAL was supported by core funding from: NIHR Blood and Transplant Research Unit in Donor Health and Genomics, UK Medical Research Council (G0800270), and British Heart Foundation (SP/09/002).

## Online Methods

### Sample collections

The ascertainment, data production, and quality control of the schizophrenia case-control whole-exome sequencing data set had been described in detail in an earlier publication^17^. Briefly, the data set was composed of schizophrenia cases recruited as part of eight collections in the UK10K sequencing project, and matched population controls from non-psychiatric arms of the UK10K project, healthy blood donors from the INTERVAL project, and five Finnish population studies. The UK10K data set was combined and analyzed with published data from a Swedish schizophrenia case-control study^24^. Sequence data for the UK10K project was deposited into the European Genome-phenome Archive (EGA) under study accession code EGAO00000000079, and the processed VCFs for the Swedish data was acquired from dbGAP under accession code (phs000473.v1.p1). The data production, quality control, and analysis of the case-control CNV data set was described in an earlier publication^26^. The schizophrenia cases were recruited as part of the CLOZUK and CardiffCOGS studies, which consisted of both schizophrenia individuals taking the antipsychotic clozapine and a general sample of cases from the UK. Matched controls were selected from four publicly available non-psychiatric data sets. All samples were genotyped using Illumina arrays, and processed and called under the same protocol. Sanger-validated *de novo* mutations identified through whole exome-sequencing in seven published studies of schizophrenia parent-proband trios were aggregated and re-annotated for enrichment analyses^13,37-42^. A full description of each trio study, including sequencing and capture technology and sample recruitment was previously described^17^.

### Sample and variant quality control

We jointly called each case data set with its nationality-matched controls, and excluded samples based on contamination, coverage, non-European ancestry, and excess relatedness^17^. A number of empirically derived filters were applied at the variant and genotype level, including filters on GATK VQSR, genotype quality, read depth, allele balance, missingness, and Hardy-Weinberg disequilibrium^17^. After variant filtering, the per-sample transition-to-transversion ratio was ~3.2 across the entire data set, as expected for populations of European ancestry^43^. For the case-control CNV analysis, we similarly excluded samples based on non-European ancestry, excess relatedness, and contamination, and only CNVs supported by more than 10 probes and greater than 10 kilobases in size were retained to ensure high quality calls. All *de novo* mutations in our study had been validated using Sanger sequencing.

We used the Ensembl Variant Effect Predictor (VEP) version 75 to annotate all variants (SNVs and CNVs) according to Gencode v.19 coding transcripts. We defined frameshift, stop gained, splice acceptor, and donor variants as loss-of-function (LoF), and missense or initiator codon variants with the recommended CADD Phred score cut-off of greater than 15 as damaging missense^29^. A gene was annotated as disrupted by a deletion if part of its coding sequence overlapped the copy number event. We more conservatively defined genes as duplicated only if the entire canonical transcript of the gene overlapped with the duplication event.

Statistical tests of the case-control exome data used case-control permutations within each population (UK, Finnish, Swedish to generate empirical *P*-values to test hypotheses. No genome-wide inflation was observed in common variant association tests and burden tests of individual genes^17^. In the curated set of *de novo* mutations, we observed the expected exome-wide number of synonymous mutations given gene mutation rates from previously validated models^30^, suggesting variant calling was generally unbiased across Gencode v.19 coding genes. Lastly, the case-control CNV data set had been previously analyzed for burden of CNVs affecting individual genes, and enrichment analyses in targeted gene sets^7,26^.

### Rare variant gene set enrichment analyses

#### Case-control enrichment burden tests

For the case-control SNV analysis, we performed permutation-based gene set burden tests using a method applied in Purcell *et al*. and implemented in PLINK/SEQ and the SMP utility^17,24^. Briefly, we first calculated one-tailed burden test-statistics for each gene:

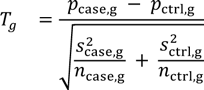

where *p_g_* is the frequency of rare variants in gene *g*, and 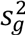 is the sample variance of gene *g*. A null distribution for *T_g_* across all genes was calculated by performing two million case-control permutations within each population (UK, Finnish, and Swedish) to control for batch and ancestry. For each gene set, an enrichment statistic was calculated as the sum of single gene burden test-statistics corrected for exome-wide differences between cases and controls:

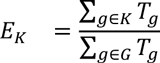

where *K* is the set of all genes in the gene set, *G* is the set of all genes in the genome, and *T_g_* is the burden statistic for gene *g*. One-sided *P*-values were calculated by comparing the observed enrichment statistic with enrichment statistics from the two million permutations. As in Purcell *et al*., the reported odds ratios and 95% confidence intervals were calculated raw counts without adjusting for case-control differences, batch and ancestry-specific differences.

#### CNV logistic regression

We adapted a logistic regression framework described in Raychaudhuri et al. and implemented in PLINK to compare the case-control differences in the rate of CNVs overlapping a specific gene set while correcting for differences in CNV size and total genes disrupted^7,18,27^. We first restricted our analyses to coding deletions and duplications, and tested for enrichment using the following model:

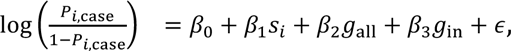

where for individual *i*, *p_i_* is the probability they have schizophrenia *i*, *s_i_* is the total length of CNVs, *g_all_* is the total number of genes overlapping CNVs, and *g_in_* is the number of genes within the gene set of interest overlapping CNVs. It has been shown that *β_1_* and *β_2_* sufficiently controlled for the genome-wide differences in the rate and size of CNVs between cases and control, while *β_3_* captured the true gene set enrichment above this background rate^7,18,27^. For each gene set, we reported the one-sided *P*-value, odds ratio, and 95% confidence interval of *β_3_*.

#### Weighted permutation-based sampling of *de novo* mutations

For each variant class of interest, we first determined the total number of *de novo* mutations observed in the 1,077 schizophrenia trios. We then generated 2 million random samples with the same number of *de novo* mutations, weighting the probability of observing a mutation in a gene by its estimated mutation rate. The baseline gene-specific mutation rates were obtained using the method described in Samocha *et al*. and adapted to produce LoF and damaging missense rates for each Gencode v.19 gene. These mutation rates adjusted for both sequence context and gene length, and were successfully applied in the primary analyses of large-scale exome sequencing of autism and severe developmental disorders with replicable results^22,31,44^. For each gene set, one-sided enrichment *P*-values were calculated as the fraction of two million random samples that had a greater or equal number of *de novo* mutations in the gene set of interest than what is observed in the 1,077 trios. The effect size of the enrichment was calculated as the ratio between the number of observed mutations in the gene set of interest and the average number of mutations in the gene set across the two million random samples. We adapted a method in Fromer *et al*. to calculate 95% credible intervals for the enrichment statistic^13^. We first generated a list of one thousand evenly spaced values between 0 and ten times the point estimate of the enrichment. For each value, the mutation rates of genes in the gene set of interest were multiplied by that amount, and 50,000 random samples of *de novo* mutations were generated using these weighted rates. The probability of observing the number of mutations in the gene set of interest given each effect size multiplier was calculated as the fraction of samples in which the number of mutations in the gene set is the same as the observed number in the 1,077 trios. We normalized the probabilities across the 1,000 values to generate a posterior distribution of the effect size, and calculated the 95% credible interval using this empirical distribution.

#### Combined joint analysis

Gene set *P*-values calculated using the case-control SNV, case-control CNV, and *de novo* data were meta-analyzed using Fisher’s combined probability method with *df* = 6 to provide a single test statistic for each gene set. We corrected for the number of gene sets tested in the discovery analysis (n = 1,776) by controlling the false discovery rate (FDR) using the Benjamini-Hochberg approach, and reported only results with a *q*-value of less than 5%.

### Description of gene sets

The full list of tested gene sets is found in Supplementary Table 1. All gene identifiers were mapped to the Gencode v.19 release, and all non-coding genes were excluded from further analysis. We first accessed and combined gene sets from five public databases: Gene Ontology (release 146; June 22, 2015 release), KEGG (July 1, 2011 release), PANTHER (May 18, 2015 release), REACTOME (March 23, 2015 release), and the Molecular Signatures Database (MSigDB) hallmark processes (version 4, March 26, 2015 release). Given our focus on very rare and *de novo* variants, we had limited power to robustly detect enrichment in small gene sets, as evident in previous studies of schizophrenia and autism rare variation in which the strongest signals came from aggregating hundreds of genes^13,24,31^. Thus, we restricted our analyses to gene sets from the five public databases with more 100 genes. We further tested a number of gene sets selected based on biological hypotheses about schizophrenia risk, and genome-wide screens investigating rare variants in broader neurodevelopmental disorders. These included gene sets described in previous enrichment analyses of schizophrenia rare variants^24^: translational targets of *FMRP*^34,45^, components of the post-synaptic density^14,24^, ion channel proteins^24^, components of the ARC, mGluR5, and NMDAR complexes^24^, proteins at cortical inhibitory synapses^7,46^, targets of mir-137^24^, and genes near schizophrenia common risk loci^11,24^. We additionally incorporated gene sets previously shown to be enriched for autism risk genes: targets of *CHD8*^31,47,48^, splice targets of RBFOX^31,49,50^, hippocampal gene expression networks^51^, and neuronal gene lists from the Gene2cognition database (http://www.genes2cognition.org)^31^. We used the pLI metric described in the ExAC v0.3 database as a measure of gene-level selective constraint^28^. Genes annotated with pLI > 0.9 were described as “highly constrained”, and those annotated with pLI < 0.9 were described as “ExAC unconstrained”. We further ranked and grouped genes into deciles and bideciles according to the pLI metric (Supplementary Figure 6 and 7). ASD risk genes were defined as genes with a FDR < 10% or < 30% by Sanders *et al*. in the largest meta-analysis of ASD exomes to date^22^. For a less stringent list, we separately defined ASD and developmental disorder *de novo* genes as genes hit by a LoF or a LoF/missense *de novo* variant in the Sanders *et al*. and the DDD study^22,44^. The DECIPHER Developmental Disorder Genotype-Phenotype (DDG2P) database (April 13, 2015 release) was used to define genes diagnostic of developmental disorders^32,44^. For a high confidence list, as used for clinical reporting in the DDD study, we included genes with a monoallelic or a X-linked dominant mode of effect and robust evidence in the literature (“Confirmed DD Genes”, “Probable DD gene”, “Both DD and IF”). From these genes, we created four lists based on mechanism (LoF or LoF/missense) and affected organ system (brain/cognition or any organ system. Finally, for background gene lists, we defined cerebellar and cortical genes as those that are expressed in at least 80% of the corresponding human brain samples in the Brainspan RNA-seq dataset^52^. A gene was defined as expressed in a sample if the exon and whole gene read counts were greater than 10 counts, and the Cufflinks lower-bound FPKM estimate was greater than 0^53^. For brain-enriched genes, we compared the differential expression of individual genes in the brain against all other tissues in the GTEx dataset^54^, and identified a subset that is 2-fold enriched with a FDR < 5%.

### Selection of allele frequency thresholds and consequence severity

To identify frequency cut-offs and functional classes that best enrich for rare variants of large effect, we looked at the magnitude and significance in two gene sets (Purcell *et al*. primary set, and Darnell *et al*. *FMRP* targets) known to be enriched for rare schizophrenia risk variants^13,24^. For schizophrenia case-control whole-exome sequencing data set, we compared enrichment at four MAF thresholds (< 0.5%, < 0.1%, singleton, and ExAC singletons), and four variant classes (synonymous, missense, damaging missense [CADD phred score > 15], and loss-of-function). The < 0.5%, < 0.1%, and singleton MAF cut-offs were defined internally in our data set. The singleton threshold restricted our analyses to variants observed only once in our data set, and the ExAC singleton threshold additionally excluded variants present in the ExAC database (v0.3). Because the ExAC v0.3 release included variants from the Swedish schizophrenia study, of which some are in our meta-analysis, we used a subset of the release containing 45,376 individuals without known neuropsychiatric disorders for the purposes of annotation. In the two gene sets, we observed no significant burden in synonymous and missense variants, with the strongest enrichment coming from LoF variants restricted to ExAC singletons (Supplementary Figure 1). The case-control signal in genes with schizophrenia *de novo* LoF and missense variants was also most significant after restricting to ExAC singletons (Supplementary Figure 1). We subsequently tested which classes of variants (LoF, LoF and damaging missense, damaging missense [CADD phred > 15], missense, and synonymous) showed the strongest enrichment in *de novo* mutations in the 1,077 parent-proband trios, and found that including both LoF and damaging missense *de novos* in a single test (P_*FMRP*_= 2.3 × 10^−5^; Pprimary = 0.033) was better powered than including just LoF (P_*FMRP*_= 0.17; P_primary_ = 0.12) and damaging missense variants (P_*FMRP*_= 2.85 × 10^−5^; P_Primary_ = 0.08). When selecting MAF cut offs for case-control CNVs, the strongest signal for known gene sets was observed when combining both deletions and duplications with a more lenient frequency cut-off (MAF < 1%, all CNVs: P_*FMRP*_ = 3.3 × 10^−5^; MAF < 0.1%, all CNVs: P_Primary_ = 0.35). However, we found that the gene set *P*-values were dramatically inflated even when testing for enrichment in the random gene sets (see below) (Supplementary Figure 2). This inflation was driven in part by very large (overlapping more than 10 genes), more common (MAF between 0.1% and 1%) CNVs observed mainly in cases or controls. Some of these, such as the known syndromic CNVs, likely harbored true risk genes. However, because these CNVs were highly recurrent in cases and depleted in controls, and disrupted a large number of genes, any gene set that included even a single gene within these CNVs would appear to be significant, even after controlling for total CNV length and genes overlapped. To ensure our model was well calibrated and its *P*-values followed a null distribution for random gene sets, we explored different frequency and size thresholds, and conservatively restricted our analysis to copy number events overlapping less than seven genes (excluding the largest 10% of CNVs) with MAF < 0.1% (Supplementary Figure 2).

### Robustness of enrichment analyses

We uniformly sampled genes from the genome (as defined by Gencode v.19) to generate random gene sets with the same size distribution as the 1,776 gene sets in our discovery analysis. For each random set, we calculated gene set *P*-values for the case-control SNV data, case-control CNV data, and *de novo* data using the appropriate method and frequency cut-offs across all variant classes. A Q-Q plot was generated using *P*-values from enrichment tests of each data set and variant type. Reassuringly, we observed null distributions in all such Q-Q plots (Supplementary Figure 4).

### Comparison of *de novo* enrichment with broader neurodevelopmental disorders

We aggregated and re-annotated de novo mutations from four studies: 1,113 severe DD probands^44^, 4,038 ASD probands^22,31^, and 2,134 control probands^23,31^. We used the Poisson exact test to calculate differences in *de novo* rates in constrained genes between SCZ, ASD, and DD and controls. Counts in each functional class (synonymous, missense, damaging missense, and LoF) were tested separately, and the one-sided *P*-value, rate ratio, and 95% CI of each comparison were reported and plotted in Figure 2, Supplementary Figure 5 and 6.

### Conditional analyses

In each of the three methods we used for gene set enrichment, we restricted all variants analyzed to those that reside in the background gene list, and tested for an excess of rare variants in genes shared between the gene set of interest (*K*) and the background list (*B*). Three brain expression gene lists based on Brainspan and GTEx, and the ExAC constrained gene list (pLI > 0.9) were used as backgrounds (see above). The enrichment statistic for the case-control SNV data was modified to:

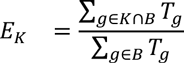

where *K* is the set of all genes in the gene set, *B* is the set of all background genes, and *T_g_* is the burden statistic for gene *g*. The logistic regression model for the case-control CNV data was modified to:

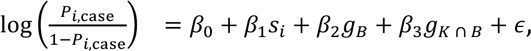

where *g_B_* is the total number of background genes overlapping a CNV, and *g*_*K* ∩ *B*_ is the number of genes in the intersection of the gene set of interest and the background list overlapping a CNV. Finally, we determined the total number of *de novo* mutations within the background gene list observed in the 1,077 schizophrenia trios, and generated 2 million random samples with the same number of *de novo* mutations. For each gene set, one-sided enrichment *P*-values were calculated as the fraction of two million random samples that had a greater or equal number of de novo mutations in genes in *K* ∩ *B* than what is observed in the 1,077 trios. Gene set *P*-values were combined using Fisher’s method. We restricted our conditional enrichment analysis to gene sets with *q*-value < 1% in the discovery analysis, and adjusted for multiple testing using Bonferroni correction.

### Rare variants and cognition in schizophrenia

The MUIR collection within the UK10K study was composed of individuals with discharge diagnoses of mild learning disability and schizophrenia (ICD-8 and -9). Evidence of remedial education was a prerequisite to inclusion, and individuals with pre-morbid IQs below 50 or above 70, severe learning disabilities, or were unable to give consent were excluded. The Schizophrenia and Affective Disorders Schedule-Lifetime version (SADS-L) in people with mild learning disability, PANSS, RDC, and DSM-III-R, and St. Louis Criterion were applied to all individuals to ensure that any diagnosis of schizophrenia was robust. More information on the recruitment guidelines in the MUIR collection can be found in a previous publication^55^.

Cognitive testing was conducted on 502 schizophrenia individuals in the UK10K data set. For these individuals, we acquired their pre-morbid IQ as extrapolated from National Adult Reading Test (NART). We retained 412 individuals for analysis after excluding all individuals with predicted pre-morbid IQ of less than 85 (or below one standard deviation of the population distribution for IQ). We additionally acquired information on educational attainment in 54 schizophrenia individuals in the UK10K dataset, and retained 27 individuals who completed at least 13 years of schooling (compulsory education in the UK). Lastly, the California Verbal Learning Test was conducted on 124 Finnish schizophrenia individuals, and a composite score was generated from measures of verbal and visual working memory, verbal abilities, visuoconstructive abilities, and processing speed. All individuals with intellectual disability were excluded from cognitive testing. Within this set of samples, we additionally excluded any individuals who ranked in the lowest decile in CVLT composite score, and retained 92 individuals for analysis. According to these criteria, we identified 531 of 697 schizophrenia individuals with cognitive data as not having intellectual disability.

Using the case-control SNV enrichment method (see above), we tested for differences in rare variant burden between the following samples: 119 SCZ individuals with ID and 6,789 matched controls, 3,614 SCZ individuals with no cognitive information and 9,343 matched controls, 531 SCZ individuals without ID and 6,789 matched controls, 119 SCZ individuals with ID and 3,614 SCZ individuals with no cognitive information, and 119 SCZ individuals with ID and 531 SCZ individuals without ID. We tested for rare variant enrichment in two gene sets: the constrained gene set (pLI > 0.9) and diagnostic DD genes with brain abnormalities as described in Decipher DDG2P database (Figure 3, Supplementary Figure 8).

Of the 1,077 probands assayed for *de novo* mutations in our meta-analysis, 619 were required to have attended mainstream schools in Bulgaria, which, at the time, excluded individuals with intellectual disability^13^. We tested for de novo enrichment in the constrained gene set (pLI > 0.9) and diagnostic DD genes with brain abnormalities, and meta-analyzed these *P*-values with the case-control SNV enrichment in 531 schizophrenia individuals without ID using Fisher’s combined probability method (*df* = 4) to provide a single test statistic.

### Consortia

#### UK10K consortium

Richard Anney, Mohammad Ayub, Anthony Bailey, Gillian Baird, Jeff Barrett, Douglas Blackwood, Patrick Bolton, Gerome Breen, David Collier, Paul Cormican, Nick Craddock, Lucy Crooks, Sarah Curran, Petr Danecek, Richard Durbin, Louise Gallagher, Jonathan Green, Hugh Gurling, Richard Holt, Chris Joyce, Ann LeCouteur, Irene Lee, Jouko Lönnqvist, Shane McCarthy, Peter McGuffin, Andrew McIntosh, Andrew McQuillin, Alison Merikangas, Anthony Monaco, Dawn Muddyman, Michael O’Donovan, Michael Owen, Aarno Palotie, Jeremy Parr, Tiina Paunio, Olli Pietilainen, Karola Rehnström, Tarjinder Singh, David Skuse, Jim Stalker, David St. Clair, Jaana Suvisaari, Hywel Williams

#### INTERVAL study

Participants in the INTERVAL randomised controlled trial were recruited with the active collaboration of NHS Blood and Transplant England (http://www.nhsbt.nhs.uk), which has supported field work and other elements of the trial. DNA extraction and genotyping was funded by the National Institute of Health Research (NIHR), the NIHR BioResource (http://bioresource.nihr.ac.uk/) and the NIHR Cambridge Biomedical Research Centre (www.cambridge-brc.org.uk). The academic coordinating centre for INTERVAL was supported by core funding from: NIHR Blood and Transplant Research Unit in Donor Health and Genomics, UK Medical Research Council (G0800270), British Heart Foundation (SP/09/002), and NIHR Research Cambridge Biomedical Research Centre.

A complete list of the investigators and contributors to the INTERVAL trial is provided in reference^56^, and http://www.intervalstudy.org.uk/about-the-study/whos-involved/interval-contributors/.

## Supplementary Tables

**Supplementary Table 1:** Full results from enrichment analyses of 1,776 gene sets. The *P*-values from enrichment tests of each variant type (case-control SNVs, DNM, and case-control CNVs) are shown for each gene set, along with the Fisher’s combined P-value (*P*_meta_) and the FDR-corrected Q-value (*Q*_meta_). N_genes_: number of genes in the gene set; SNV: single nucleotide variants from whole-exome data; DNM: *de novo* mutations.

**Supplementary Table 2:** Gene sets enriched for rare coding variants conferring risk for schizophrenia at FDR < 5%. The effect sizes and corresponding *P*-values from enrichment tests of each variant type (case-control SNVs, DNM, and case-control CNVs) are shown for each gene set, along with the Fisher’s combined *P*-value (*P*_meta_) and the FDR-corrected Q-value (*Q*_meta_). N_genes_: number of genes in the gene set; Est: effect size estimate and its lower and upper bound assuming a 95% CI; SNV: single nucleotide variants from whole-exome data; DNM: *de novo* mutations.

**Supplementary Table 3:** Results from enrichment analyses of FDR < 1% gene sets, conditional on brain-expressed and ExAC constrained genes. We restrict enrichment analyses to genes that reside in four different background gene sets, three defined on brain expression (Cerebellum and cortex-expressed genes in Brainspan, brain-enriched expression in GTeX) and the fourth on genic constraint (ExAC-constrained genes), and determined if gene sets with FDR < 1% in the meta-analysis still had significance above the specific background. The *P*-values from enrichment tests of each variant type (case-control SNVs, DNM, and case-control CNVs) are shown for each gene set, along with the Fisher’s combined P-value (*P*_meta_). N_genes_: number of genes in the gene set; SNV: single nucleotide variants from whole-exome data; DNM: *de novo* mutations.

